# Call patterns encode and transmit emotion in marmoset monkeys

**DOI:** 10.1101/2022.08.03.502601

**Authors:** Junfeng Huang, Hailin Liu, Chen Wang, He Ma, Yongkang Sun, Liangtang Chang, Neng Gong

**Author notes:** Correspondence to (N.G.). These authors contributed equally (Junfeng Huang, Hailin Liu).

## Abstract

Marmoset monkeys have attracted much attention as a non-human primate model for studying vocal communication, but the call pattern and its meaning in marmoset communication are largely unknown. Here, we analyze sounds produced by hundreds of marmosets either in isolation or in pairs and reveal distinct call patterns in marmoset communication. The most prominent phee calls could be categorized into multiple grades based on the number of comprising phee syllables. Call transitions exhibited non-random patterns, favoring transition to the same or adjacent grade, with long sequences limited within two adjacent grades. The interval, composition, and temporal distribution of calls were significantly different between isolated and paired marmosets. Notably, different patterns of phee calls correlated with the heart rates and emotional states of marmoset, with the higher call grade reflecting a more agitated state. Antiphonal calling also exhibited distinct patterns and phee calls directly affected the heart rate of the listener in a manner depending on the grade of phee calls. Thus, phee call patterns in marmosets could encode emotional states and transmit emotion between turn-taking marmosets. How emotional expression in animals evolves into semantic communication in humans remains a mystery. Such complex call patterns in marmoset vocalization could represent the evolutionary prelude to semantic communication in primates.

## Main text

Common marmoset (*Callithrix jacchus*), a New World monkey species, has attracted much attention as a useful model for studying social behaviors, especially vocal communication and monogamous cooperative breeding ^1–3^. Marmoset monkeys have a complex vocal repertoire consisting of rich call types both in captivity and natural environment ^4,5^. They show typical vocal turn-taking and produce trill and phee calls for close and long-distance communication, respectively^6,7^. Marmoset vocalizations also exhibit a certain degree of plasticity from infancy to adulthood ^8^. Calls made by infant marmosets in the family group undergo dynamic developmental change in a manner that depends on both physical maturation and contingent parental feedback ^9,10^. The influence of parental feedback on vocal development was also found in a juvenile stage (3 postnatal months) ^11,12^. Adult marmosets can precisely control vocal behavior by rapidly interrupting ongoing vocalizations ^13^, and are also able to adaptively modify their vocal structure when they encountered interfering sounds ^14^. These rich call types and flexible call structures could enable the emergence of complex vocal communication in marmosets. However, exactly how marmosets use different call patterns to encode and transmit information remain unknown. In this study, we have delineated the organizational rules of marmoset call sequences, and demonstrated that call patterns could be used for communicating the emotional state of the marmoset.

## Results

### Categorization of various grades of phee calls

When kept in isolation, marmosets produce mostly phee calls, which are known to be used for long-distance vocal communication between conspecifics ^7^. In this study, we recorded vocalizations of 110 adult marmosets (54 males and 56 females; age 6.1 ± 0.3 years; weight 390 ± 7 g, SEM), each for 30 - 90 min in isolation (see Methods, Fig. 1A). For a total of 28,622 discrete phee syllables acquired (Fig. 1B), we measured inter-syllable intervals (ISIs) between all adjacent syllables (n = 28,512 events) and found two distinct groups of ISIs (Fig. 1C), as indicated by clear bimodal pattern in ISI distribution that fitted well with two independent Gaussian distributions. Two distinct peaks of the ISI distribution profile were found at 0.32 s and 10.67 s (Fig. 1D), consistent with a previous finding ^15^. Such bimodal pattern was largely invariant among marmosets, as shown by the heat plot of ISI distributions for all 110 animals (Fig. 1E). Examples of ISI distributions for three marmosets (#27, #57, #75; Fig. 1, F to H) showed that each marmoset exhibited two peak values of ISIs that were slightly variable. These peak values for all 110 marmosets were clearly separated into two groups with the median at 0.35 s and 10.00 s (Fig. 1I), close to that found for pooled phee syllables pooled from all marmosets (Fig. 1D).

**Fig. 1.**
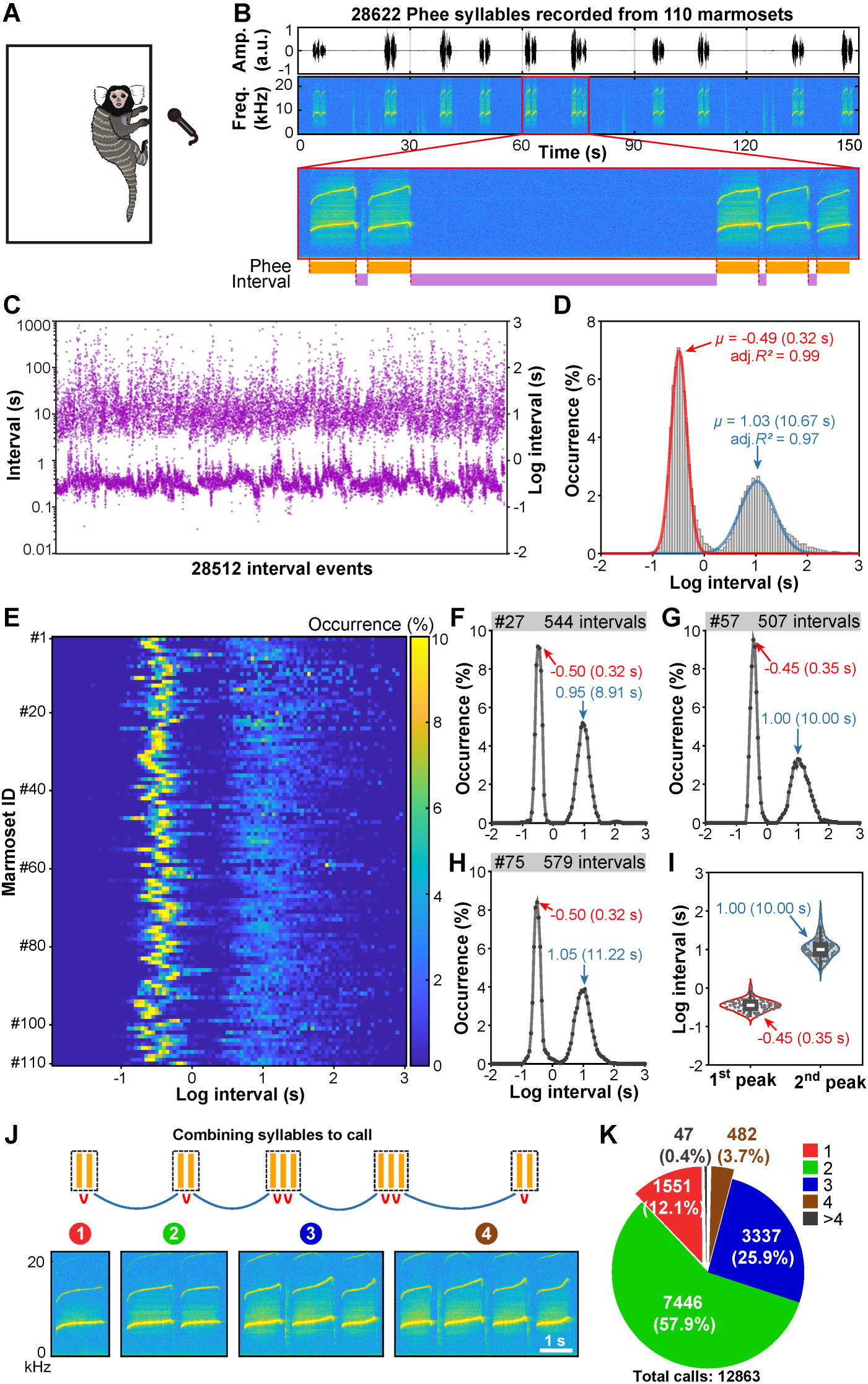
Categorization of various grades of phee calls. (**A**) An illustration of the call recording method in which a marmoset was isolated in a cage in the sound-proof room and recorded for 30 - 60 minutes. (**B**) A typical example showing the recorded phee syllable sequence and the measurement of inter-syllable intervals (ISIs). (**C**) Plot of all ISIs in a log scale. (**D**) Distribution of ISIs in log scale and the fitting curves of Gaussian functions. (**E**) The heat plot of ISI distributions for all 110 animals. (**F** to **H**) Examples of ISI distributions for three marmosets #27, #57, and #75. The peaks were showed with arrows. (**I**) The first peak and second peak of each marmoset’s ISIs distribution (gray dot). The white lines indicated the median values of these two peaks in 110 marmosets, which were 0.35 s and 10.00 s respectively. (**J**) According to a 1-s threshold, closely spaced phee syllables were grouped together into different phee calls. (**K**) Pie diagram showing the proportion of each call grade.

The interval of 1 s (log value = 0) could well separate ISIs into two groups of short and long ISIs (Fig. 1J). We then defined closely spaced syllables with short intervals to be a call (or a “word”), and long intervals represent intervals between calls. All 28,622 phee syllables recorded were thus grouped into 12,863 phee calls, which could be further categorized into calls of different grades by the number of syllables contained in the call, with N-phee representing phee call containing N syllables. We found that 2- phee was the most common (57.9%), followed by 3-phee (25.9%), 1-phee (12.1%), and 4-phee (3.7%) (Fig. 1K). Calls containing more than 4 syllables were rare (∼0.4%, Fig. 1K), thus were excluded in the following analyses.

### Organization rules of phee call transitions

To examine the organization of phee call sequences (Fig. 2A for 3 example marmosets; Extended Data Fig. 1 for 110 marmosets), we used a matrix to display the frequency of call transitions to the same or a different grade in the call sequences from all 110 marmosets (Fig. 2B), and transition probabilities, as depicted by the thickness of arrow bars, are shown in Fig. 2C. We found that the percentage for transitions into a call of the same grade (“Rep”: repetition, 66.3%) was about two-fold of that for transition to another grade (“Trans”, 33.7%). Most (92%) of the latter was one-grade transition (Trans-One) that occurred between calls of adjacent grades, and the rest was skip-grade transition (Trans-Skip, 8%, Fig. 2C). Furthermore, the percentages of Trans-Up (transition to a higher grade, T_AB_) and Trans-Down (transition to a lower grade, T_BA_) were largely symmetrical (Fig. 2D). Analysis of the inter-call intervals for all repetitions and transitions showed that the interval to the next call showed a similar transition-direction dependence for all call grades (Extended Data Fig. 2). Overall, the inter-call interval for repetition (11.2 s, median; n = 8408) was shorter than that of Trans-Up (8.7 s, median; n = 2170, *P* < 0.001), and longer than that of Trans-Down (15.9 s, median; n = 2096, *P* < 0.001) (Fig. 2E). Thus, the ordering of marmoset phee calls follows distinct rules that prefer repetition and one-grade transition, with transition direction-dependent inter-call intervals.

**Fig. 2.**
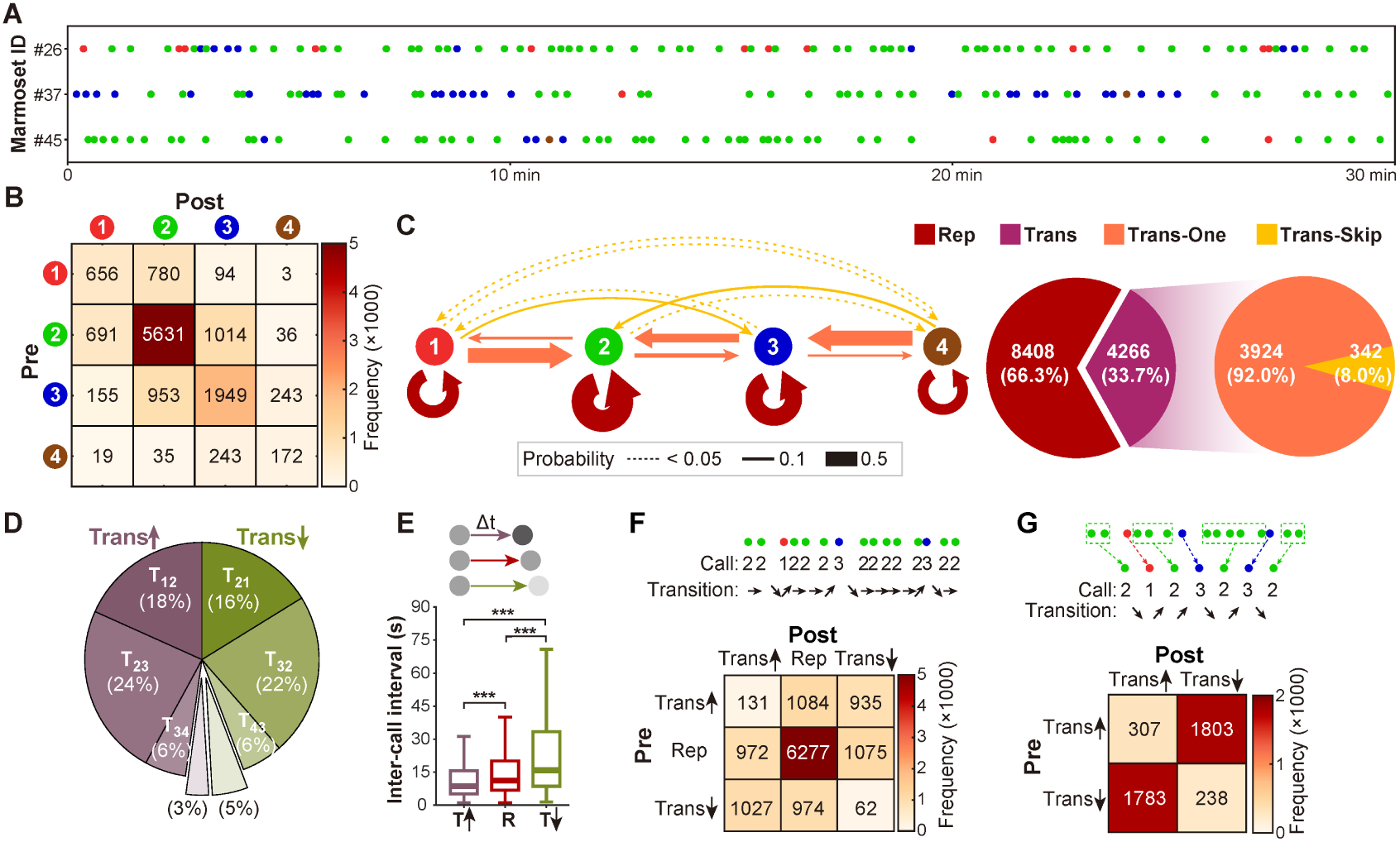
Organization rules of phee call transitions. (**A**) Three examples showing the phee call sequences of the marmosets ^#^26, ^#^37, and ^#^45. (**B**) Matrix showing the frequency of transitions between two sequential calls including data from all 110 marmosets. (**C**) Transition diagrams for call sequences. Arrow line color and thickness represented the type and the probability of transition, respectively. Pie diagram showing the occurrence percentage of each transition type. “Rep”: repetition; “Trans”: transition to a call of different grade; Trans-One: one-grade “Trans”; Trans-Skip: skip-grade “Trans”. (**D**) Pie diagram showing the occurrence percentage of each Trans-Up (transition to a call of higher grade, left) or Trans-Down (transition to a call of lower grade, right). T_AB_: transition from call A to call B. (**E**) Box diagram showing inter-call intervals for transition types that inter-call intervals were shorter in Trans-Up, longer in Trans-Down than that of repetitions (Median; \*\*\**P* < 0.001, linear mixed model). T: “Trans”; R: “Rep”. (**F**) One example of call sequence showed transition information (“Rep”, Trans-Up and Trans-Down), and the relationship between transition types from 110 marmosets was shown in frequency matrix. (**G**) Sequential repeated calls were grouped into a single “call” and the frequency matrix contained only Trans-Up and Trans-Down after omitting “Rep”.

The existence of a long-range sequencing rule for call transitions was further studied by examining the influence of the particular type of call transition on the occurrence of subsequent transitions. We first analyzed the frequency of various patterns of two sequential transitions, as shown by the frequency matrix in Fig. 2F. We found that sequential repetitions (“Rep” to “Rep”) occurred with the highest frequency, and Trans-Up or Trans-Down tended to be followed by “Rep” or a “Trans” of the opposite direction (Fig. 2F). The long-range relationship between Trans-Up and Trans- Down transitions in a call sequence was further revealed by omitting the “Rep” between two “Trans” transitions, via grouping of sequential repeated calls into a single call (Fig. 2G). The resulting frequency matrix containing only Trans-Up and Trans-Down (Fig. 2G) showed that two sequential “Trans” transitions were much more likely to be opposite in direction. These results demonstrate the existence of long-range rule, namely, long sequences of transitions were largely limited within two adjacent call grades. Overall, sequential and long-range transitions in marmoset calls tended to avoid large changes in call grade.

### Call patterns during vocal turn-taking

Following the studies of phee call sequences in isolation, we further examined whether the organization rules for call sequences were altered during vocal turn-taking. Using 25 adult marmosets couples (age 4.5 ± 0.2 years; weight 403 ± 8 g; SEM, n = 50), we recorded vocalizations (for ∼ 60 min) of each marmoset of the couple separated by a curtain (Fig. 3A, see Methods). First, we analyzed sequencing rules of each marmoset’s own vocalizations during turn-taking. A total of 9905 phee syllables were acquired, and syllable density during vocal turn-taking (3.3 ± 0.3 per min, SEM, n = 50) was significantly lower than that in isolation (4.3 ± 0.3 per min, SEM, n = 110, *P* = 0.019, Mann–Whitney U test). The inter-syllable intervals during vocal turn-taking also distributed into two distinct groups, as shown by the heat plot of ISI distribution (Extended Data Fig. 3), but intra-call syllable intervals (first peak) were shorter and inter-call intervals (second peak) were longer as compared to those found in isolation (Fig. 3B). By the same criterion of 1-s threshold, all phee syllables were grouped into 4836 phee calls (call sequences shown in Extended Data Fig. 4), and the call composition was markedly modified by vocal turn-taking (Fig. 3C). This modification was also observed in each individual, as shown by the averaged proportion of 1-phee significantly elevated from 13.1 ± 1.6 % (SEM, n = 110) to 23.0 ± 3.4% (SEM, n = 50, P = 0.017, Mann–Whitney U test). By analyzing the transition patterns, we found that the call ordering, transition probabilities, and long-range effects were nearly identical to that observed in isolation (Extended Data Fig. 5). The shortening of intra- call syllable intervals was also similar across different call grades (Fig. 3D). The prolonged inter-call intervals still depended on the transition direction: Comparing to that for repetitions (15.5 s, median; n = 3116), the intervals were shorter for Trans-Up (12.5 s, median; n =822, *P*< 0.001) and longer for Trans-Down (24.1 s, median; n = 797, *P* < 0.001) (Fig. 3E and Extended Data Fig. 6). These results indicate that sequence rules of phee calls during vocal turn-taking were generally similar to those found in isolation, but the intra- and inter-call intervals as well as call compositions were significantly modified.

**Fig. 3.**
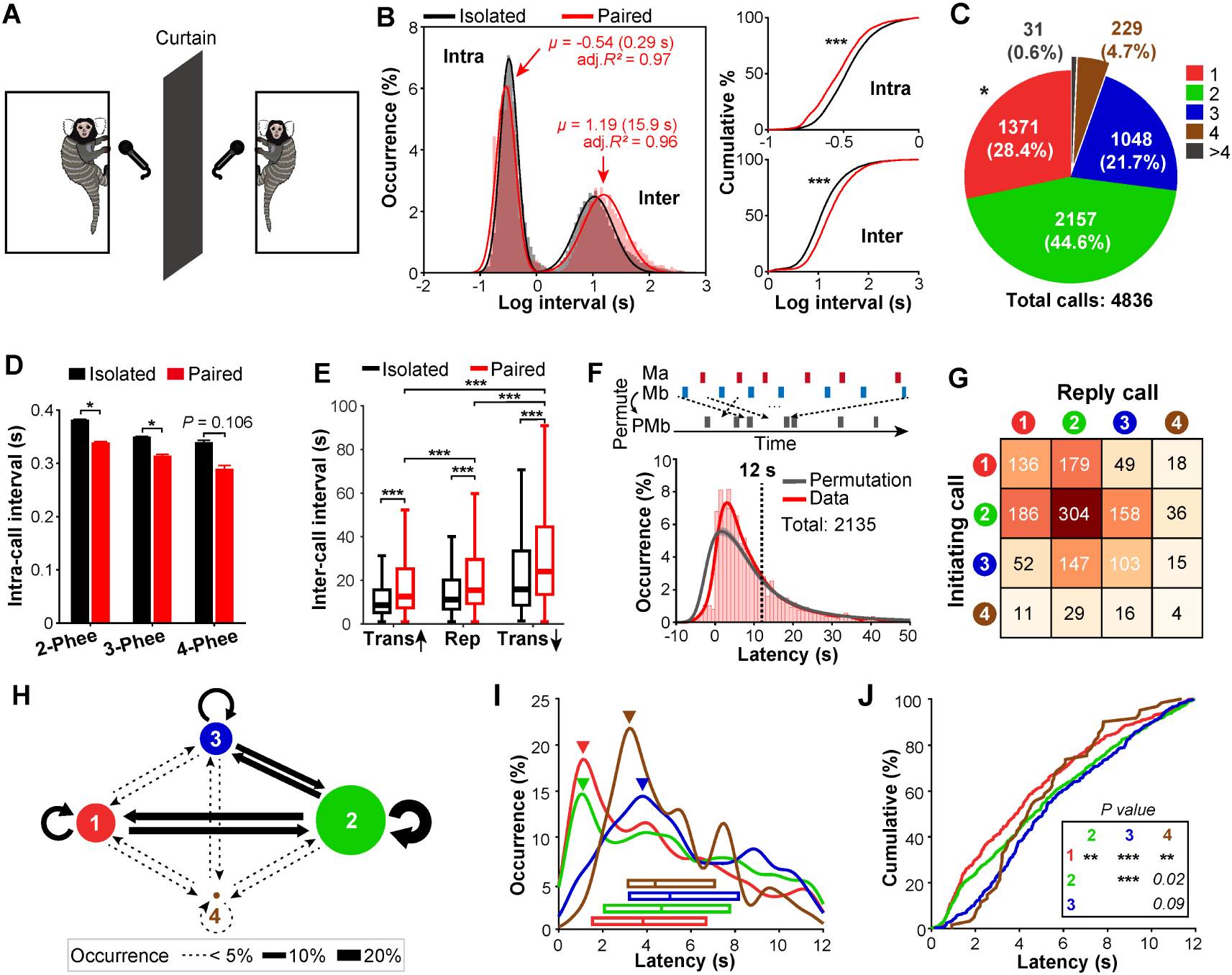
Call sequences during vocal turn-taking. (**A**) An illustration of the vocal recording of two marmosets in pairs. (**B**) Distributions of inter-syllable intervals in log scale and Gaussian fitting curves of both isolated and paired conditions. Right: cumulative distributions of intra-call syllable intervals and inter-call intervals (\*\*\**P* < 0.001, Kolmogorov-Smirnov test). (**C**) Pie diagram showing the proportion of each call grade in paired condition (\**P* < 0.05, Mann-Whitney U test, n = 160 marmosets). (**D**) For each call grade, intra-call syllable intervals were shorter in pairs than those in isolation (Mean ± SEM; \**P* < 0.05, linear mixed model). (**E**) Inter-call intervals of different transition types in isolation and in pairs (Median; \*\*\**P* < 0.001, linear mixed model). (**F**) Distribution of reply latencies between two sequential calls made by two marmosets for both original and randomly permuted sequences. A call pair with a latency less than 12 s (dash line) was considered as an antiphonal call pair. Top panel: an illustration of permutation by randomly shuffling the sequence of inter-call intervals. Bottom panel: distributions of reply latencies of experimental data (red) and permuted data (gray, Mean with 95% confidence interval). (**G**) Frequency matrix showing the call grades of antiphonal call pairs. (**H**) Reply diagrams of antiphonal call pairs. Diameter of the solid circle represented the proportion of each call grade. Arrow line thickness represented the occurrence percentage of each call pair. (**I**) Distribution and median of the reply latency which was elicited by the initiating call of each grade. Arrow indicated the peak of occurrence percentage. Red: 1-phee, green: 2-phee, blue: 3-phee, brown: 4-phee. (**J**) Cumulative distribution of the reply latency which was elicited by the initiating call of each grade. (\*\**P* < 0.01, \*\*\**P* < 0.001, Kolmogorov-Smirnov test).

We then examined the rules for antiphonal calling during vocal turn-taking. Two sequential calls made by different monkeys was considered to be a call pair in communications. We found a total of 2135 call pairs and analyzed the distribution of reply latency in each pair (Fig. 3F). We confirmed the presence of vocal turn-taking by showing that the distribution of latencies was distinctly different from that found after random permutation of marmosets’ vocal sequences in each pair (Fig. 3F). According to the distribution of latencies, a call pair with a reply latency less than 12 s was considered as an effective antiphonal call pair, similar to that used previously ^16^. Among 2135 call pairs, a total of 1443 effective antiphonal call pairs were observed, and the frequency (Fig. 3G) and occurrence percentage (Fig. 3H) for each type of antiphonal call pair showed that marmosets preferred to use 1-phee and 2-phee for antiphonal calling. Furthermore, the reply call tended to be in the same or adjacent but not non-adjacent grade relative to the grade of the initiating call (Fig. 3H). The latency for each initiating call eliciting a reply call of any grade was found to be significantly different for 4 grades of initiating calls, with 1-phee eliciting the more rapid reply than other grades of initiating calls (Fig. 3, I and J, see original data in Extended Data Fig. 7). This supports the notion that shorter latency in turn-taking facilitates marmoset vocal communication ^6,16^. These results revealed the rule of antiphonal calling during vocal turn-taking in marmosets, suggesting that different phee call patterns could convey information reflecting isolated vs. paired conditions.

### Call patterns encode marmosets’ emotional states

To address potential information conveyed by phee calls, we first analyzed the temporal distribution of different phee calls in isolation, and found that high-grade phee calls (containing more phee syllables) appeared earlier (Fig. 4A). Given that marmosets appeared nervous when entering a new environment, different phee calls may be associated with the emotional state. We also recorded the tsik call, which is known to be a specific vocal index of anxiety and fear in marmosets ^17^, in isolation and during vocal turn-taking (Fig. 4B) and found that tsik calls mainly occurred in the early stage of recording (Fig. 4B), further supporting the notion that marmosets were in a nervous state when they entered the recording room. By analyzing the correlation of the timing of phee and tsik calls, we found that the number of tsik occurred near (within 5 s before and after) the phee call was significantly correlated with the phee call grade (Fig. 4C) – the number of tsik calls associated with 4- and 3-phee was significantly higher than that with 1- and 2-phee (Fig. 4C). Moreover, for 2-, 3- and 4-phee, the number of tsik calls following the phee call was significantly higher than that before the phee call, whereas no such difference was found for 1-phee calls (Fig. 4D). Thus, different grades of phee calls correlate with marmoset’s emotional state, with the higher call grade reflecting a more anxious state.

**Fig. 4.**
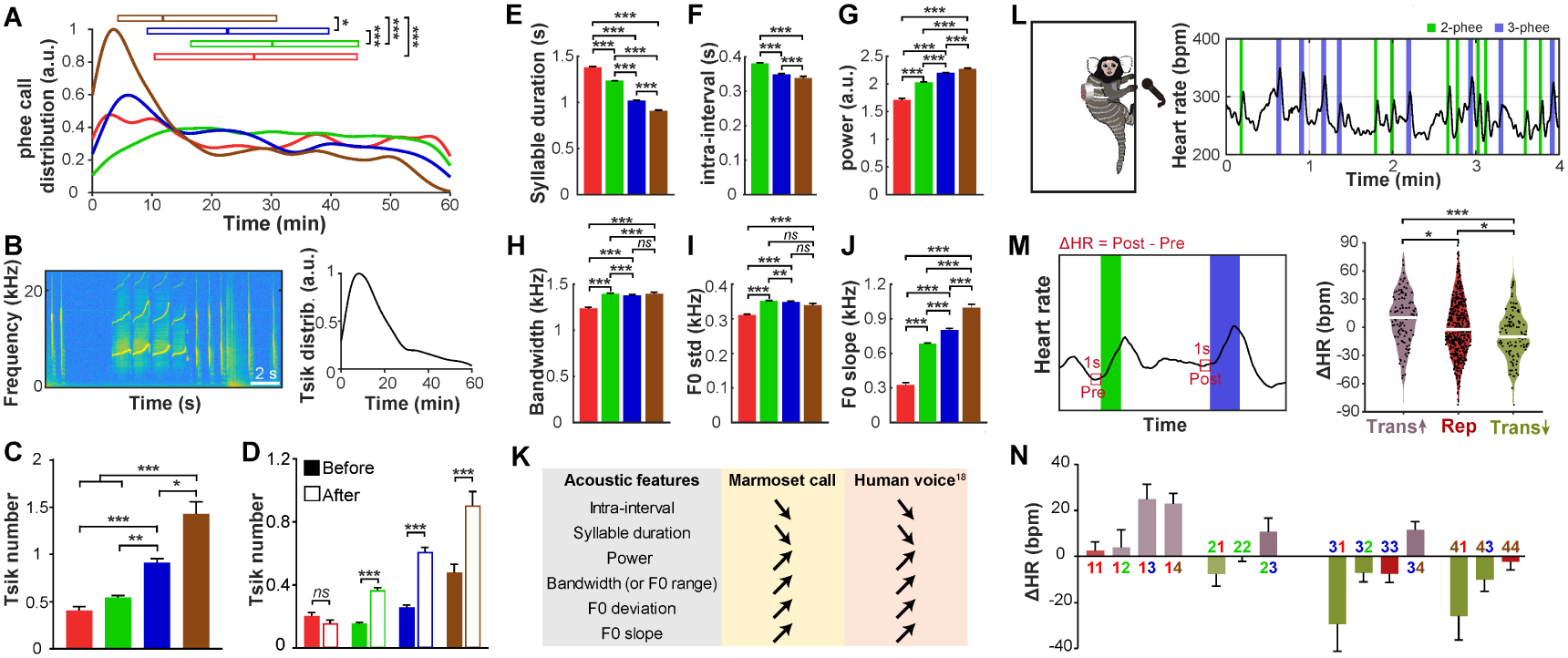
Call patterns encode emotion of marmoset. (**A**) Distributions and medians of the starting time of calls with different grades in isolation. The distributions were normalized by the max value of all distribution curves (n = 87 marmosets, \**P* < 0.05, \*\*\**P* < 0.001, Kruskal-Wallis test). Red: 1-phee, green: 2-phee, blue: 3-phee, brown: 4-phee. (**B**) Spectrogram of the exemplar tsik calls (left) and temporal distribution of tsik calls (right, n=87 marmosets). (**C**) The number of tsik calls that were either 5 s before or after the phee call, which were shown respectively in (**D**) (\**P* < 0.05, \*\**P* < 0.01, \*\*\**P* < 0.001, linear mixed model). (**E** ∼ **J**) The acoustic features of phee syllables, including syllable duration, intra-call interval, syllable power, frequency bandwidth, standard deviation of fundamental frequency (F0) and the slope of F0. (**K**) Acoustic features of marmoset high-grade calls resembled that of human in emotional state associated with high arousal. The human acoustic data were modified from ref. *18*. (**L**) An illustration of vocalization and electrocardiogram recording in isolation (left), and an example piece of marmoset’s heart rates during vocalizing (right). (**M**) Change of heart rate in call transition. Left: an illustration of calculating change of heart rate (ΔHR), which was calculated as the averaged 1-s heart rate before the post call minus the averaged 1-s heart rate of the pre call. Right: change of heart rate in Trans-up, Rep and Trans-down (n = 6 marmosets and white lines indicated medians, \**P* < 0.05, \*\*\**P* < 0.001, linear mixed model). (**N**) ΔHR of all specific transition types (Mean ± SEM). The Bar color and the number above/beneath the bar showed the type of call transition.

In human speech, the acoustic features could reflect the emotional state associated with high arousal ^18^. Many acoustic features of marmoset phee calls were distinctly different among phee syllables of various call grades, consistent with a previous finding^19^. We found that syllables of higher-grade calls showed shorter duration (Fig. 4E) and intervals (Fig. 4F), higher power (Fig. 4G) and bandwidth (Fig. 4H), as well as higher deviation (Fig. 4I) and slope (Fig. 4J) of fundamental frequency. Such call grade-dependent trends in acoustic features of phee syllables were similar to that found for the acoustic features of human sounds produced under increasing arousal (Fig. 4K). We also examined the heart rate change as another physiological index for the emotional state of the marmoset. We found that marmosets’ heart rates consistently showed a rapid increase during the production of phee calls (Fig. 4L). By examining the average heart rate during 1-s period before each call, presumably reflecting the emotional state prior to call production, we found that the direction of heart rate changes was the same as that of call transitions (Fig. 4M) –increased for Trans-Up, unchanged for Rep, and decreased for Trans-Down, as shown by the distribution of heart rate changes (Fig. 4M). The same trend of heart rate change was also be observed when data for each particular type of call transitions were analyzed (Fig. 4N). Thus, the acoustic features and sequence patterns of phee calls reflect the emotional state of marmosets.

### Call patterns transmit emotion between paired marmosets

To explore the possibility of emotion transmission between the turn-taking marmosets, we examined whether call sequence patterns of the caller marmoset affect that of the other marmoset. First, by comparing to that found for marmoset in isolation, the temporal distribution of phee calls of marmosets in pairs was significantly changed, especially for 1-phee (Fig. 5A), which became much more frequent during the late half of the recording period (Fig. 5B). The antiphonal call pair initiated by 1-phee and replied by 1-phee occurred also more frequently in the late stage of recording (Fig. 5, C to E), indicating that the paired animals used 1-phee increasingly more often with time during turn-taking. Furthermore, the occurrence of tsik call near (5 s before and after) the phee call was again highly correlated with the phee call grade, with 1-phee having the fewest tsik calls around it (Fig. 5F). Thus, in the presence of communication, the paired marmosets appeared to transmit the emotion, leading to temporal increase of 1-phee calls of the pair. Finally, we further test this idea by examining whether the heart rate of the listener is affected by the caller’s call grade and found that the occurrence of phee calls immediately affected the listener’s heart rate in a manner depending on the grade of caller’s phee calls (Fig. 5G). Consistent with the emotional state carried by each phee, 1-phee significantly decreased the listener’s heart rate and 2-phee had no significant effect, while 3- and 4-phee significantly increased the listener’s heart rate (Fig. 5G). Thus, our data provide direct evidence that call patterns can transmit emotion in marmoset communication.

**Fig. 5.**
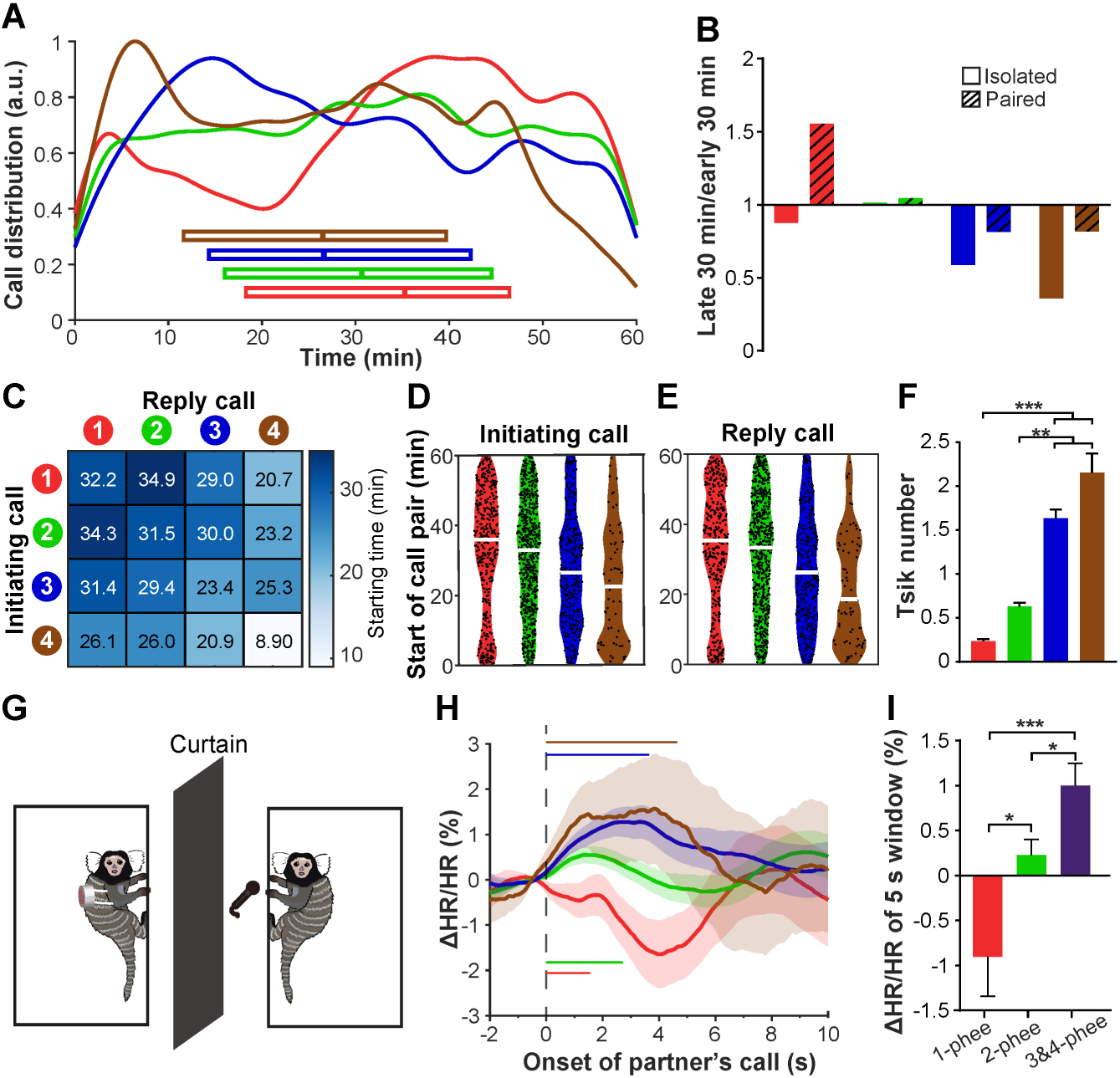
Call patterns transmit emotion between paired marmosets. (**A**) Distributions and medians of the starting time of calls with different grades in paired condition (n = 50 marmosets). The distributions were normalized by the max value of all distribution curves. Red: 1-phee, Green: 2-phee, Blue: 3-phee, Brown: 4- phee. (**B**) The ratio of phee call occurrences in the late recoding period (30 - 60 min after the start of recording) to that in the early recording period (0 - 30 min after the start of recording) in isolation and in pairs. (**C** ∼ **E**) The starting time of call pair in vocal turn-taking. In (**C**), the matrix showed the time when call pair started, with the number indicating the median starting time of call pair of each type. The starting time of call pair which was initiated or replied by specific phee call was shown in (**D**) and (**E**), respectively. (**F**) The number of tsik calls that was either 5 s before or after the phee call in paired condition. (**G**) An illustration of vocalization and electrocardiogram recording in paired condition. (**H**) The heart rate response (ΔHR/HR_0_, see Methods) when hearing different phee calls, which were aligned by the call onset (n = 13 marmosets, mean ± SEM; the horizontal lines indicated the average duration of different phee calls and the dash line indicated the call onset). (**I**) The average change of heart rate in 5 s window after the call onset (Mean ± SEM, \**P* < 0.05, \*\*\**P* < 0.001, ANOVA test).

## Discussion

By recording of vocalizations from hundreds of marmosets either in isolation or in pair, we performed detailed analyses of their vocal sequence patterns and uncovered distinct rules in syllable combinations and call transitions. Sequence patterns during vocal turn-taking generally followed those observed in isolation, but intra-call syllable intervals, inter-call intervals, and call compositions were significantly modified. These suggests that the call pattern of marmosets is probably determined by both intrinsic constraints in vocal production and external influences due to social interactions. Many songbird species could produce songs that have complex syntax with syllable sequences following certain rules ^20–23^, which are flexible under different contexts ^24^. Although there are some similarities, the sequencing rules we observed in marmosets are quite different from those found in birds. The call sequence in marmosets we examined are simply the combination and ordering of the same phee syllable, whereas bird songs contain multiple syllables. Furthermore, unlike that in bird songs, phee call intervals of marmosets are much longer than the word intervals in human language. On the other hand, we found that distinct phee call patterns in marmosets could encode emotional states and transmit the emotion between turn-taking marmosets.

It is well known that acoustic features can reflect emotional changes in both animals and humans. Here we found that the emotional states of marmosets could be reflected in both call patterns and the acoustic features of the call. Our results indicate that phee calls comprising various numbers of syllables could reflect different emotional states, with calls of rapid repetitive syllables indicating a more agitated state. The direction of call transitions also reflects the tendency of emotional changes. Inter-call intervals depended on the direction of call transition, suggesting that the propensity for the marmoset to become more agitated than to calm down. Sequential and long-range call transitions tended to maintain calls within two adjacent grades, implicating drastic changes of emotional states are unlikely. Despite the difficulty in accurately characterizing emotional states, we obtained both vocal and physiological evidence that different patterns of phee calls could transmit emotion among marmosets. The reply latency is relatively long during marmoset vocal turn-taking, perhaps reflecting the time required in emotional contagion. Vocal and locomotor coordination in marmosets is associated with the activity of autonomic nervous system ^25^, which is also involved in emotion processing. Furthermore, some neurons in marmoset’s amygdala are found to specifically respond to phee calls, supporting the idea that phee calls convey emotional information. Our findings call for further study of the neural mechanism by which phee call patterns encode and transmit emotional states in marmosets.

Ever since Darwin ^26^, animal vocalization has been thought to be evolved for expressing emotion. Human language is composed of ordered sequences of syllables, which convey meaning by forming words and sentences. To draw a distinct boundary between the evolution of semantic communication and emotion expression has been difficult ^27^. Together with the evolution of the vocal production apparatus ^28^, the emergence of call sequence patterns in non-human primates represents an evolutionary basis for increasing complexity of vocal communication in primates. The transition in the function of vocal patterns from representing emotion to conveying semantic meaning could be an important process for the evolution of human language.

## Methods

### Subjects

In this study, we recorded vocalizations of 110 adult common marmosets (54 males and 56 females; aged 1.5 - 13.5 years and weighted 220 - 625 g) in isolated condition and 25 pairs of marmoset couples (25 males and 25 females; aged 1.9 - 9.5 years and weighted 308 - 516 g) in paired condition. In addition, another 6 marmosets (2 males and 4 females; aged 2.2 – 7.6 years) and 10 pairs of marmoset couples (10 males and 10 females; aged 1.7 – 8.4 years) were used for heartbeat recording in isolated condition and in paired condition, respectively. All marmosets were captive and lived in family groups. Each marmoset family was housed in a wire-mesh cage (L × W × H: 0.90 m × 0.80 m × 0.85 m), equipped with a sleeping box and other enrichment materials. Marmosets had ad libitum access to food and water, and the breeding rooms were maintained at a temperature range from 26°C to 30°C and a 12-h: 12-h light-dark cycle. Animal care and experimental procedures were approved by the Animal Care and Use Committee at the Institute of Neuroscience, Center for Excellence in Brain Science and Intelligence Technology, Chinese Academy of Sciences.

### Experimental setup and vocal recording

All vocal recordings were conducted between 9:00 and 18:00 in a soundproof recording room (L × W × H: 4.5 m × 1.5 m × 2.2 m) from 2013 to 2023. For the recording in isolated condition, each marmoset was transported by a transfer box from the breeding room to a wire-mesh cage (L × W × H: 0.90 m × 0.80 m × 0.85 m) at the height of 1.20 m above the floor in the recording room. A directional microphone (Audio-Technica AT2031) was set directly in front of the cage at a distance of 0.35 m, and the audio signal was digitized through an audio interface (Icon Utrack or Roland Octa-Capture UA-1010) at 44.1or 48 kHz sample rate with 16 bits by Adobe Audition 3.0 software. Each marmoset was recorded for 30 - 90 min with the mean duration of 61.6 min. 87 marmosets, whose recording durations were longer than 60 min, were used to analyze temporal distributions of phee calls. In paired condition, two marmosets of a couple were removed to the recording room and placed separately in two cages about 3.0 m apart, with a directional microphone set directly 0.35 m in front of each cage. An opaque black curtain was hung between the cages to occlude visual contact between marmosets. Each couple was recorded for 60 - 74 min with the mean duration of 60.7 min.

### Vocalization detection and classification

The audio signals were digitally high-pass filtered at 3 kHz, and a custom-made Matlab script was used to detect the onset and offset of each syllable by applying an energy threshold. Then, the types of vocalizations (mainly phee calls and tisk calls) were annotated manually in Praat software based on their spectrograms. The detection and classification results were cross-checked by 2 - 3 researchers. In both isolated and paired conditions, recorded vocalizations were almost phee calls, the specific long-distance contact call in marmosets. In total, we recorded 28,622 and 9,905 phee syllables in the isolated and paired condition, respectively.

### Vocalization analysis

#### Distribution of inter-syllable interval

Inter-syllable interval (ISI) was defined as the silent period between two sequential phee syllables. After plotting the ISIs in a logarithmic (log10) scale, the ISI distribution displayed a bimodal pattern with two peaks, each mode of which was fitted with one Gaussian function:

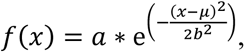

where *x* is the interval and *f(x)* is the occurrence probability. For each individual marmoset, the ISI distribution was smoothed by averaging the value of three nearby bins in the histogram (Fig. 1, F to H), and two peak ISI values in the distribution were measured. All these two peak values for 110 marmosets were plotted as two groups and their distributions were smoothed with a normal kernel function in Fig. 1I. According to these distributions, a 1 s interval was set as the threshold to separate ISIs into two groups of short and long ISIs.

#### Ordinal sequence of phee calls

During recording, marmosets vocalized multiple phee calls which formed call sequences (Extended Data Fig. 1). To examine the organization of phee call sequences, the frequency of bigrams in call or call transition, namely two sequential phee calls (Fig. 2B) or transitions (Fig. 2F), were counted. For further analysis of transition direction, the “Repetition” was removed by combining the calls with same grades, and then the remained transitions were only “Trans Up” or “Trans Down” (Fig. 2G).

#### Time window of antiphonal calling

Vocal turn-taking of phee calls was popular in communication between two marmosets. To analyze the call sequence in paired condition, two sequential phee calls vocalized by different marmosets were considered to be a call pair. The duration of interval between the offset of the initiating call and the onset of the reply call was regarded as reply latency, with negative value representing overlapping of two calls. To determine the time window of antiphonal calling in turn-taking, we did a permutation test as follows. First, for each marmoset, we permuted its partner’s phee call sequence by shuffling the order of inter-call intervals randomly. This permutation was performed 1000 times to generate a permuted data set. Second, the distribution of the reply latencies was calculated. Last, we compared the distribution of the real data (2,135 call pairs) with that of the permuted data set, and calculated the time window during which the distribution of the real data was higher than 97.5% percentile of the distribution of the permuted data. The time window of antiphonal calling was determined as 0 - 12 s.

#### Acoustic features of phee calls

Since the phee call might had multiple syllables, we calculated the acoustic features of each syllable at first. The audio recordings were resampled at 44.1kHz and the phee syllables were labeled manually as shown above. Then the audio recordings were processed with a Butterworth bandpass filter from 2kHz to 20kHz to eliminate background noise. The acoustic features of each syllable were calculated as follows. Syllable duration is the time duration between the onset and offset of the syllable; Intra-interval is the interval between the offset of the preceding syllable and the onset of the following syllable. Power was calculated as the integral of the square of the syllable amplitude per second; Bandwidth was the range between the lowest and highest frequencies of the syllable, the spectrum amplitudes of which were 5% of the maximum amplitude of the spectrum; To calculate the fundamental frequency (F0), the audio signal was processed with a Butterworth bandpass filter from 4 kHz to 14 kHz and was framed into short-time frames by a 512-point Hanning window with 50% overlapping and the F0 trace was the curve of the frequency with maximum energy in each frame. The standard deviation of F0 was the standard deviation of F0 over all frames; The slope of F0 was calculated as the difference of F0 between syllable onset and offset and then was normalized by the syllable duration. For phee call with multiple syllables, each acoustic feature was averaged across the comprising syllables and this mean acoustic feature was considered as the call feature.

### Electrocardiogram recording

The subject was anesthetized with 3% isoflurane to remove hair on the chest and on the back by shaving and applying hair removal cream (Veet) 3- 5 days before ECG recording. On the day of recording, a train of square waves were delivered by a data acquisition card (National Instruments, USB-6351) to an ECG recorder (Quanlan, LanMao X7minni) and an audio interface (Roland, Octa-Capture UA-1010) simultaneously as the first trigger signal to synchronizing ECG data and audio data. Then the subject was transported from the colony room to the recording room and the portable ECG recorder was put on the back of the subject with one electrode on the left chest and another electrode on the left back. The ECG recorder was tied by adhesive tape properly to make sure good signal-to-noise ratio (SNR) and to avoided interfering marmoset’s movement and vocalization. The ECG signal was digitized at 500Hz and saved at the recorder in real time. The marmoset was placed in the cage in the recording room for 60 minutes, and after that, the ECG recorder was removed from the marmoset and the second synchronization trigger signal was delivered to the ECG recorder and the audio interface. We recorded vocalizations and ECG for 60 min from 6 marmosets and 10 pairs of marmosets in isolated or paired condition, respectively. In paired condition, 7 marmosets did not vocalize so the ECG data of their partners were not included in further analysis and the other 13 marmosets used for analysis were 8 males and 5 females with their ages ranging from 1.9 – 8.4 years.

### Heart rate analysis

For analysis, the ECG signal was first processed with a Butterworth bandpass filter from 10 Hz to 100 Hz to remove direct-current component and potential jitters. Then the peaks of R waves of the signal were labeled and extracted by an open-source Matlab script ^29^, which were further checked and adjusted by 3 experimenters. Overall, the SNR of most data was good enough to extract the peaks of R waves accurately, but several small clips of the data were discarded since its low SNR. The heart rate was calculated as the reciprocal of the interval between the peaks of two adjacent R waves. The heart rates were smoothed by applying a moving average filter with 10 data points.

In isolated condition, to analyze the relationship between call transition and hear rate, we compared the difference of the average heart rate of 1-s period before vocalizing between two adjacent phee calls (ΔHR = HR of 1-s period before the following phee call – HR of 1-s period before the preceding phee call). In paired condition, we analyzed that how the phee call of one marmoset affected the heart rate of its partner. In detail, we extracted the heart rate of the listener marmoset and interpolated the data linearly with a 0.01-s time bin. The averaged heart rate of 1-s period before the onset of the phee call was set as baseline heart rate (HR0) and we calculated the heart rate response to the phee call as: ΔHR/HR0 = (HR -HR0)/HR0, where HR was the heart rate and the HR0 was the baseline heart rate. This normalization method could show the change of heart rate and it was also applied popularly in data processing of other areas such as fluorescence response in calcium imaging ^30^. According to the ΔHR/HR0 traces, we set a time window from 0 s to 5 s after the onset of the phee call as the response window and compared the ΔHR/HR0 in this window among different call grades.

### Statistical analysis

All statistical analyses were accomplished with SPSS Version 21.0 and all statistical tests were two-tailed. In our recordings, marmoset vocalized various number of phee calls which were not fully independent of each other, so linear mixed model (LMM) was conducted to compare intra-call syllable intervals, inter-call intervals, tisk number, acoustic features and the change of heart rate among groups with marmoset identity as random effect. The intra-call syllable intervals, inter-call intervals and syllable power were log-transformed (log10) to meet the normality assumption of LMM and Bonferroni corrections were applied for all post-hoc pairwise comparisons. Kolmogorov-Smirnov test was conducted to compare different cumulative distributions. Mann-Whitney U test was conducted to compare the phee syllable density and the proportion of phee calls between isolated condition and paired condition. To compared the temporal distributions of phee calls with different grades, we calculated the averaged starting time of phee call for each grade in each marmoset and then performed a Kruskal-Wallis test with Bonferroni corrections for pairwise comparisons. An ANOVA test with Games–Howell post hoc test was applied to compare the heart rate response among different call grades (3-phee and 4-phee were combined together because of their small number).

## Data availability

The data that support the findings of this study are available from the corresponding author upon request.

## Code availability

MATLAB scripts used in our data analysis are available from the corresponding author upon request.

## Acknowledgments

We thank Mu-ming Poo, Liping Wang and Zhen-Hua Ling for helpful discussions and insightful comments on this manuscript, Yishan Xie, Ruixin An, Yuan Xin, Xiangrui Li, Xinyi Yu and the marmoset facility at CAS Center for Excellence in Brain Science and Intelligence Technology for assistance in sound recordings, heartbeat measurements and analyzing. This work was supported by Science and Technology Innovation 2030-“Brain Science and Brain-inspired Research” Major Project Grant No. 2021ZD0203900, Shanghai Municipal Science and Technology Grant No. 22ZR1481500, NSFC Project 32371085, “Strategic Priority Research Program” of the Chinese Academy of Sciences, Grant No. XDB32010000, Shanghai Municipal Science and Technology Major Project, Grant No. 2018SHZDZX05, CAS Key Technology Talent Program to N.G., Shanghai Yangfan Program, No. 22YF1461100 and China Postdoctoral Science Foundation, No. 2023M731485.

## Author contributions

N.G. conceived and designed the study. J.H., H.L., C.W., H.M., Y.S. and L.C. performed sound recordings. J.H. and C.W. performed heartbeat measurements. J.H. H.L., and H.M. analyzed data and prepared the figures. N.G. and J.H. wrote the manuscript. All authors have read and approved the manuscript submission.

## Materials & Correspondence

Correspondence and requests for materials should be addressed to N.G.

