## Supplemental Information for "Call patterns encode and transmit emotion in marmoset monkeys"

### Extended data:

### Extended Data Figure 1

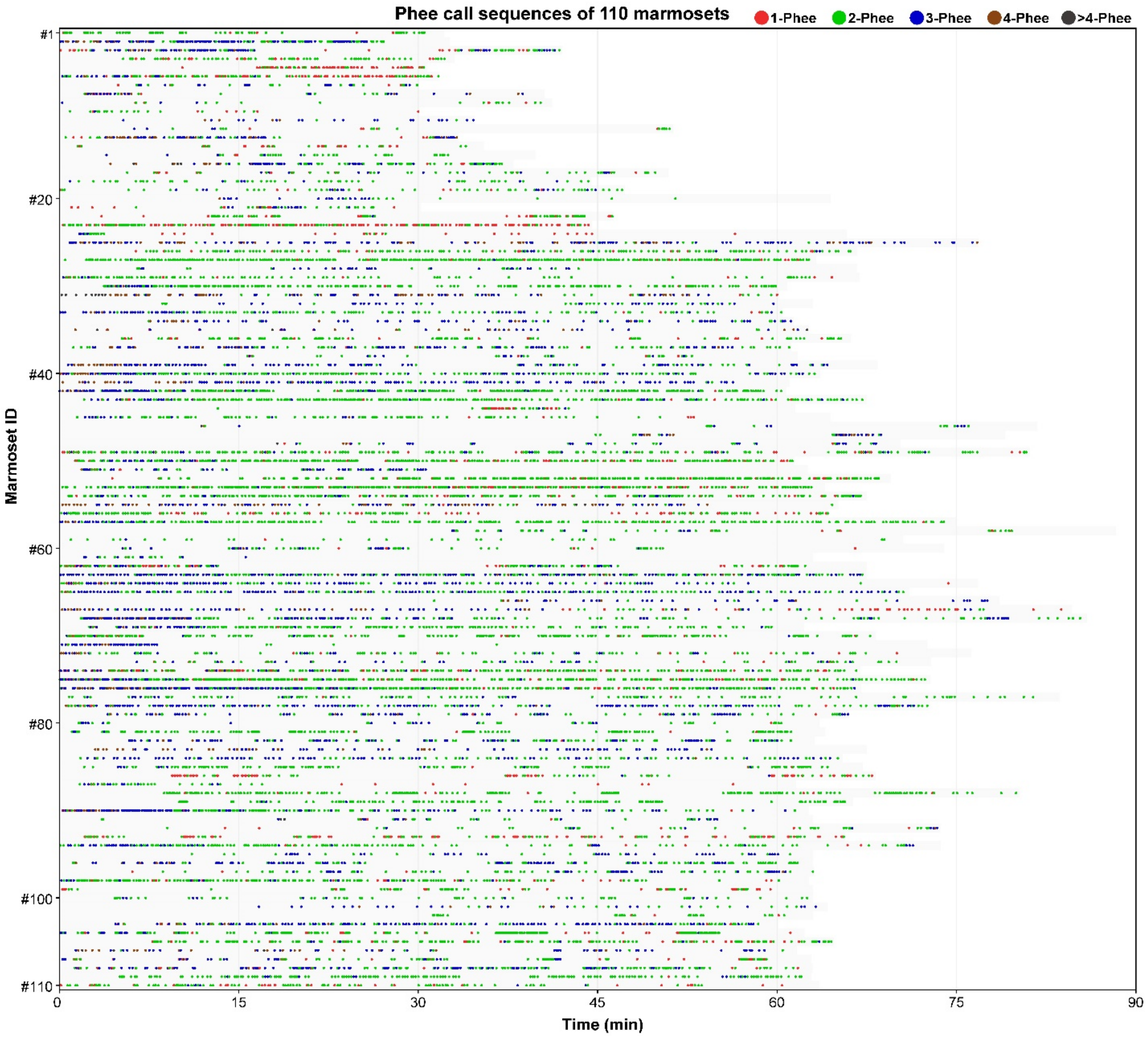


**Extended Data Fig. 1. Phee call sequences of all 110 marmosets in isolation.**

Each row represented one marmoset. Different phee calls were showed as different color dot. Red: 1-phee, green: 2-phee, blue: 3-phee, brown: 4-phee, dark gray: > 4-phee. The light gray shadow in each marmoset indicated the recording duration.

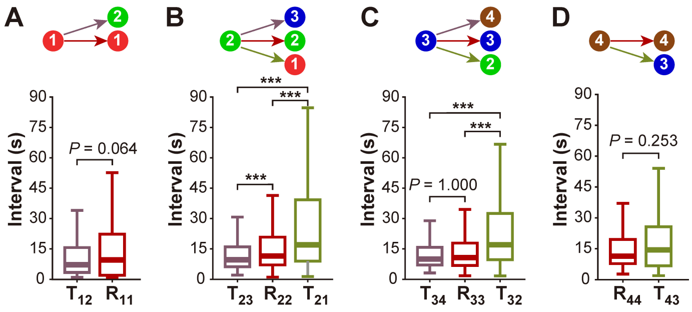


**Extended Data Fig. 2. The inter-call intervals for all specific transition types in isolation.**

(**A~D**) The inter-call intervals between two sequential phee calls, which were initiated by 1-phee, 2-phee, 3-phee and 4-phee, respectively (Median; ***$P < 0.001$, LMM). T: "Trans"; R: "Rep".

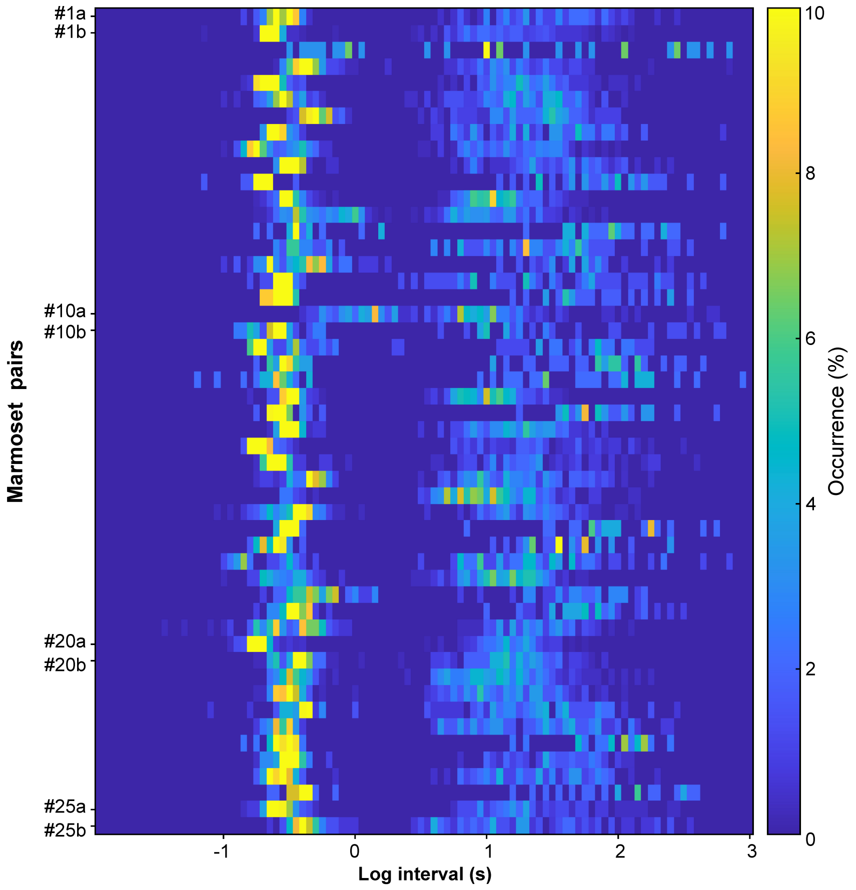


**Extended Data Fig. 3. The heat plot of inter-syllable interval distributions for all 25 marmoset pairs in pairs.**

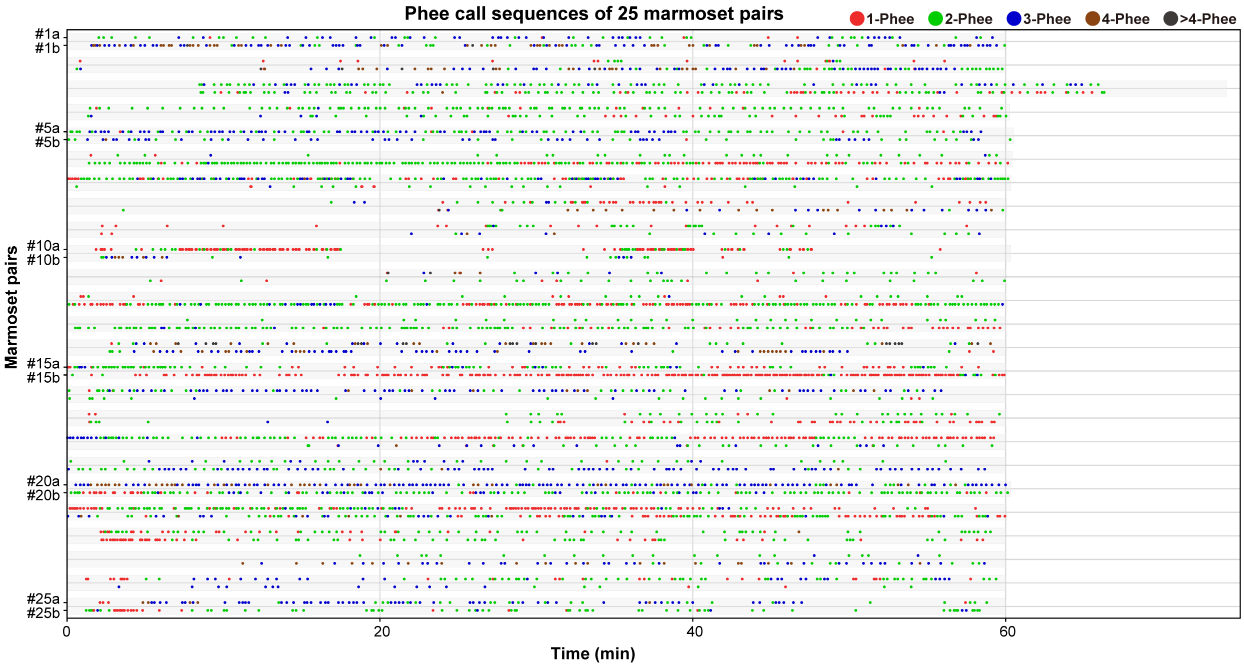


**Extended Data Fig. 4. Phee call sequences of all 25 marmoset pairs in paired condition.**

Each row represented one marmoset. Different phee calls were showed as different color dot. Red: 1-phee, green: 2-phee, blue: 3-phee, brown: 4-phee, dark gray: > 4-phee. The light gray shadow in each marmoset indicated the recording duration.

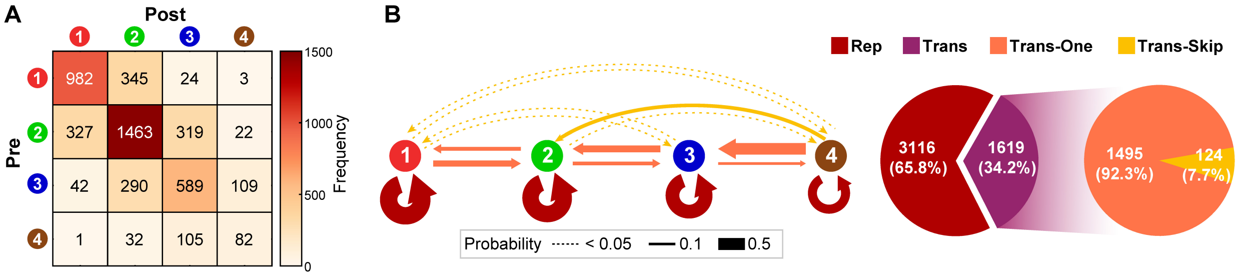


**Extended Data Fig. 5. Ordinal rules of sequential phee calls in each marmoset's own vocalizations in paired condition.**

(**A**) Matrix showing the frequency of all transitions between two sequential calls including data from all 25 marmoset pairs. (**B**) Transition diagrams for call sequences. Arrow line color and thickness represents the type and the probability of transition, respectively. Pie diagram showing the occurrence percentage of each transition type. "Rep": repetition; "Trans": transition to a call of different grade; Trans-One: one-grade "Trans"; Trans-Skip: skip-grade "Trans".

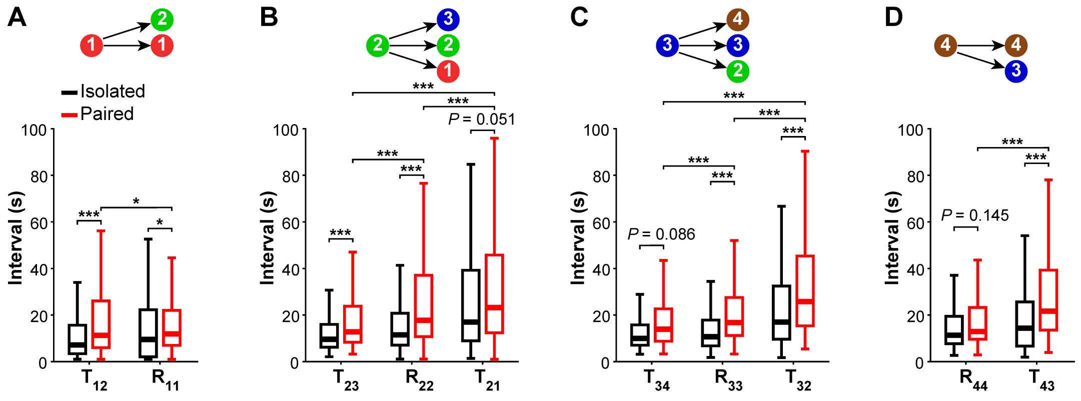


**Extended Data Fig. 6. Inter-call intervals were prolonged in paired condition and remained dependent on the transition direction.**

(**A ~ D**) Box diagrams showing inter-call intervals for all specific transition types both in isolation and in pairs (Median; **P* < 0.05, ****P* < 0.001, LMM).

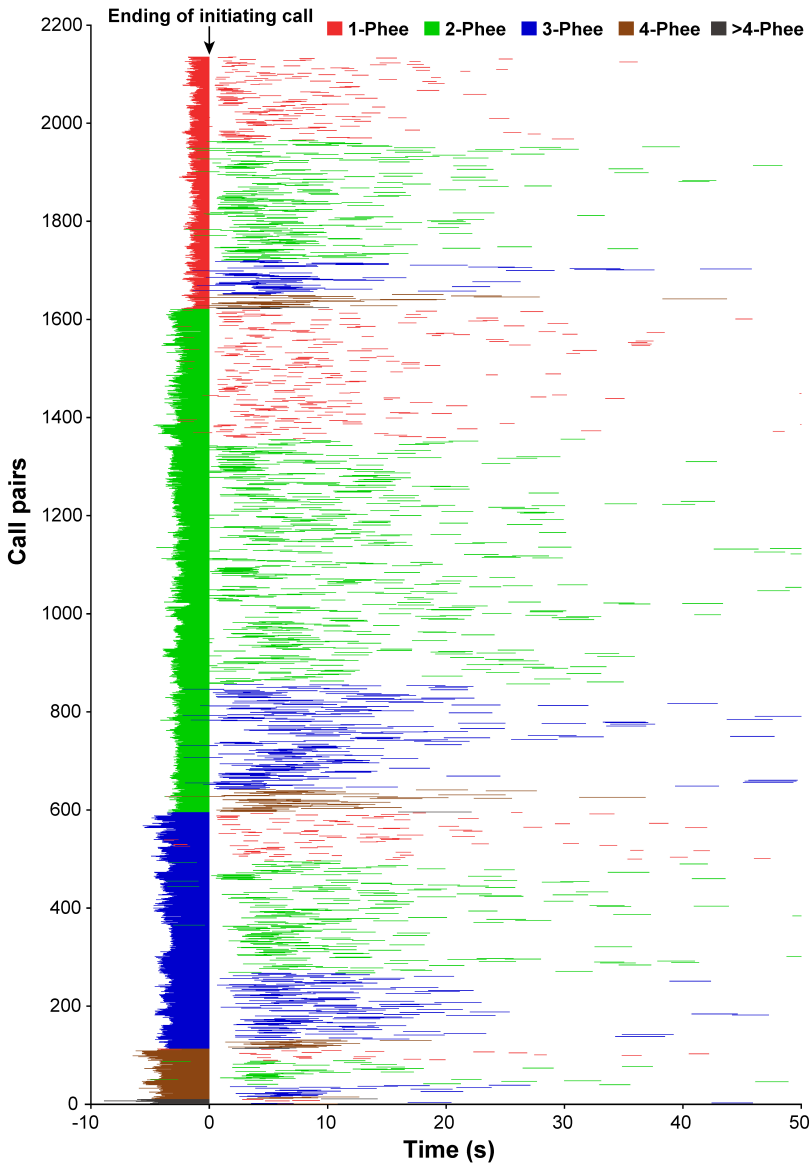


**Extended Data Fig. 7. All call pairs produced by two marmosets in pairs.**

Each row showed one call pair and the length of the color line represented the duration of phee call. The ending of the initiating call was aligned to the zero point of time.
